# Dynamic urine proteome changes in a rat model of simvastatin-induced skeletal muscle injury

**DOI:** 10.1101/2021.06.10.447866

**Authors:** Jing Wei, Yuhang Huan, Ziqi Heng, Chenyang Zhao, Youhe Gao

**Affiliations:** Department of Biochemistry and Molecular Biology, Beijing Normal University, Gene Engineering Drug and Biotechnology Beijing Key Laboratory, Beijing 100875, China

**Author notes:** These authors contributed equally to this work (Jing Wei, Yuhang Huan). Corresponding author: Youhe Gao.

**Keywords:** Urine proteome, statin-associated muscle symptoms, animal model, biomarkers

## Abstract

**Background:** Statin-associated muscle symptoms (SAMS) are the main side effects of statins. Currently, there are no effective biomarkers for accurate clinical diagnosis. Urine is not subject to homeostatic control and therefore accumulates early changes, making it an ideal biomarker source. We therefore examined urine proteome changes associated with SAMS in an animal model.

**Methods:** Here, we established a SAMS rat model by intragastric intubation with simvastatin (80 mg/kg). Biochemical analyses and hematoxylin and eosin (H&E) staining were used to evaluate the degree of muscle injury. The urine proteome on days 3, 6, 9 and 14 was profiled using liquid chromatography coupled with tandem mass spectrometry (LC-MS/MS) with the data-independent acquisition (DIA) method.

**Results:** Differential proteins on day 14 of SAMS were mainly associated with glycolysis/gluconeogenesis, pyruvate metabolism, metabolism of reactive oxygen species and apoptosis, all of which were reported to be associated with the pathological mechanism of SAMS. Among the 14 differentially expressed proteins on day 3, FIBG, OSTP and CRP were associated with muscle damage, while EHD1, CUBN and FINC were associated with the pathogenic mechanisms of SAMS. MYG and PRVA increased dramatically compared with CK elevation in serum on day 14 of SAMS.

**Conclusions:** Our preliminary results indicated that the urine proteome can reflect early changes in the SAMS rat model, providing the potential for monitoring drug side effects in future clinical research.

## Introduction

Statin-associated muscle symptoms (SAMS), which are the main side effects of statins, are reported in 10% to 25% of patients receiving statin therapy. The defined syndromes included symmetrical muscle pain, weakness and/or accompanying creatine kinase (CK) elevations >3× the upper limit of normal (ULN)[1]. According to muscle damage severity, SAMS is usually divided into myalgia, myopathy, myositis and rhabdomyolysis[2]. The clinical diagnosis of SAMS usually involves the combination of clinical assessment and biochemical indicator detection, such as the patient’s self-description of muscle symptoms[1], the evaluation of muscle symptoms after suspended statin administration[3], the measurement of creatine kinase (CK) levels after statin administration, the assessment of the statin myalgia clinical index (SMCI)[1] and the evaluation of skeletal muscle by 31-phosphorus magnetic resonance spectroscopy[4]. Although there are currently many ways to potentially diagnose SAMS, it is still very difficult to perform an accurate clinical diagnosis of SAMS. The first reason is that the diagnosis of SAMS is subjective to some extent and may be associated with the patients’ own health, their self-assessment or other drugs[5]. It is essential for clinicians to distinguish SAMS from other types of muscle pain[6]. Second, it is time-consuming to evaluate whether muscle symptoms will be relieved by ceasing statin therapy, which may also increase the heart burden of patients. Finally, effective biomarkers to diagnose SAMS are still lacking. CK levels are not specific enough, as they can still increase after regular exercise[7]. In addition, skeletal troponin I (sTnI), myosin light chain 3 (Myl3), creatine kinase M isoform (Ckm), and fatty acid binding protein 3 (Fabp3) were reported to be valuable for the diagnosis of drug-induced skeletal muscle injury; however, as muscle injury was induced by various drugs, the specificity and clinical transformation values of these proteins are still under assessment[8]. Therefore, there is an urgent need to find novel biomarkers that can detect SAMS early and sensitively, which can not only save time for the assessment of statin intolerance but can also provide important support for the adjustment of appropriate medications in time.

As a rapidly developed analytical tool, mass spectrometry-based urinary proteomics is designed to establish a novel, noninvasive liquid biopsy diagnostic method, which is used in the detection of various clinical diseases[9,10]. However, as urine is easily affected by various external factors, such as sex, diet, age and medications[11], it is still challenging to find direct associations with related diseases. Animal models are an effective way to solve this problem, as they can not only minimize these external factors but also allow for the ability to monitor the whole process of disease development, making it possible to detect early changes even before pathology changes and clinical manifestations are present. In fact, the urine proteome of animal models has already been used to detect early biomarkers before any related pathological changes occur in tumor models such as lung tumors[12], liver tumors[13], and brain tumors[14]; organ fibrosis models such as liver fibrosis[15] and lung fibrosis[16]; and organ inflammation models such as chronic pancreatitis[17] and autoimmune myocarditis[18]. Urine has also been reported to be a sensitive biomarker source, as it is more sensitive than blood[19] and can reflect related changes even with limited tumor cells[20]. Despite these valuable merits of the urine proteome, however, the urine proteome has not been applied in predicting the side effects of drugs. In this research, we aimed to determine whether the urine proteome can reflect changes associated with SAMS.

It has been reported that simvastatin is the formulation most frequently associated with adverse muscle effects in the clinical setting[21]. In addition, simvastatin is also the most commonly used statin in rodent models of SAMS[2,22]. In this study, the SAMS rat model was established by intragastric intubation with simvastatin using a stainless curved feeding needle lasting for 14 days, as reported previously[23]. Urine samples were collected on days 3, 6, 9, and 14. Biochemical analyses and hematoxylin and eosin (H&E) staining were used to evaluate the degree of skeletal muscle injury. The dynamic urinary proteomes were profiled using liquid chromatography coupled with tandem mass spectrometry (LC-MS/MS) with the data-independent acquisition (DIA) method. This study was designed to identify early urine proteome changes associated with SAMS. The study design flowchart is presented in Figure 1.

**Figure 1.**
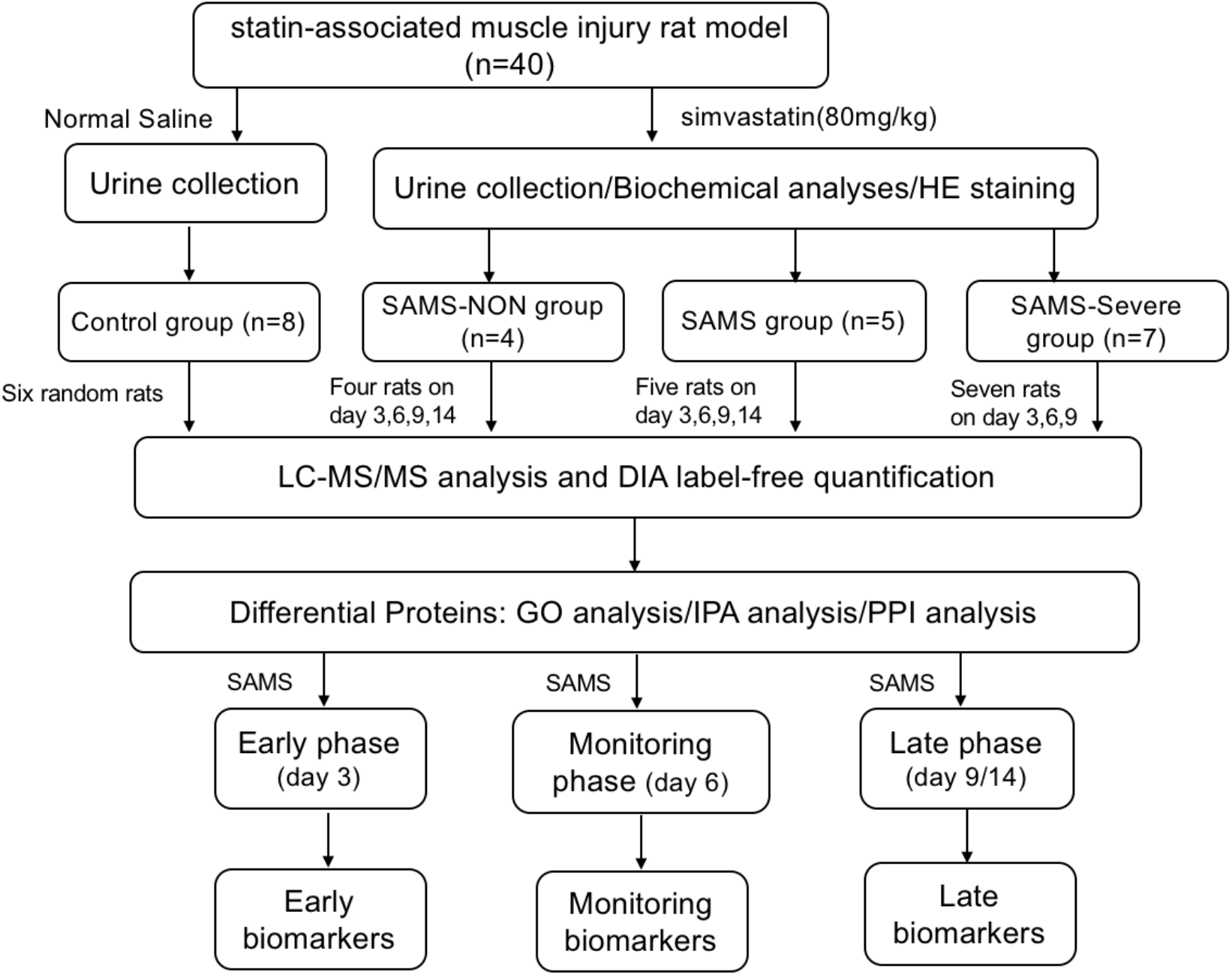
Technical flowchart of urinary protein identification in statin-associated muscle injury rats. Urine samples were collected, extracted, digested, and identified by liquid chromatography coupled with tandem mass spectrometry (LC-MS/MS) identification. Functional analysis of differential proteins was performed by GO, IPA and PPI. Candidate biomarkers of muscle injury in different phases were also identified.

## Materials and Methods

### 1. Animals and treatment protocols

Female Wistar rats (n=40, 200 ± 20 g) were purchased from Beijing Vital River Laboratory Animal Technology Co., Ltd. All animals were housed with a standard laboratory diet under controlled indoor temperature (21 ± 2°C), humidity (65–70%) and 12-h/12-h light–dark cycle conditions. The experiment was approved by the ethics and animal welfare committee of Beijing Normal University (Approval ID: CLS-EAW-2020-032). All experiments were performed in accordance with relevant guidelines and regulations of the National Health Commission and the Ministry of Science and Technology.

Simvastatin was purchased from Merck, Sharp and Dohme (West Point, PA, USA) as 40-mg tablets. Forty rats were randomly divided into two groups: the control group (n=8) and the simvastatin-treated group (n=32). The simvastatin-induced muscle injury rat model was established as previously described. In the simvastatin-treated group, rats received simvastatin (80 mg/kg per day in saline, 2 ml) by intragastric intubation using a stainless curved feeding needle for 14 days. In the control group, rats received an equal volume of NS.

### 2. Histopathology

Four rats in the simvastatin-treated group in the control group were randomly sacrificed on days 3, 6, 9 and 14 using an overdose of sodium pentobarbital anesthetic. The control group were all sacrificed on day 14. Muscle tissues, including the gastrocnemius (GAS) and soleus (SOL), were quickly fixed in 10% formalin. Then, the samples were embedded in paraffin, sectioned, and evaluated with hematoxylin and eosin (H&E) staining.

### 3. Biochemical analyses

Blood samples were collected from the abdominal vena cava on days 3, 6, 9 and 14 in the simvastatin-treated group. The blood samples in control group were collected on day 14. Then, serum was separated at 3000 rpm for 10 min within 2 h of collection and stored at −20°C for analysis within one week. The serum concentrations of alanine aminotransferase (ALT), aspartate aminotransferase (AST), blood urea nitrogen (BUN), cholesterol (CHO), triglyceride (TG), high-density lipoprotein (HDL), low-density lipoprotein (LDL) and creatine kinase (CK) were measured to assess the degree of liver damage, kidney damage and skeletal muscle damage.

### 4. Urine sample preparation

Before urine collection, all rats were housed in metabolic cages for 2–3 days. Urine samples were collected from the control group and the simvastatin-treated group on days 3, 6, 9 and 14. All rats were placed in metabolic cages individually for 12 h to collect urine without any treatment (at least 6 mL). After collection, the urine samples were centrifuged at 3000 × *g* for 30 min at 4°C and then stored at −80°C.

For urinary protein extraction, the urine samples were first centrifuged at 12,000 × *g* for 30 min at 4°C. Then, 4 mL of urine from each sample was precipitated with three volumes of ethanol at −20°C overnight. The pellets were dissolved in lysis buffer (8 mol/L urea, 2 mol/L thiourea, 50 mmol/L Tris, and 25 mmol/L DTT). Finally, the supernatants were quantified by the Bradford assay.

A total of 100 μg of protein was digested with trypsin (Trypsin Gold, Mass Spec Grade, Promega, Fitchburg, WI, USA) using FASP methods [24]. The protein in each sample was loaded into a 10-kDa filter device (Pall, Port Washington, NY, USA). After washing two times with urea buffer (UA, 8 mol/L urea, 0.1 mol/L Tris-HCl, pH 8.5) and 25 mmol/L NH4HCO3 solutions, the protein samples were reduced with 20 mmol/L dithiothreitol (DTT, Sigma) at 37°C for 1 h and alkylated with 50 mmol/L iodoacetamide (IAA, Sigma) for 45 min in the dark. The samples were then washed with UA and NH4HCO3 and digested with trypsin (enzyme-to-protein ratio of 1:50) at 37°C for 14 h. The digested peptides were desalted using Oasis HLB cartridges (Waters, Milford, MA, USA) and then dried by vacuum evaporation (Thermo Fisher Scientific, Bremen, Germany).

### 5. Spin column separation

The digested peptides were dissolved in 0.1% formic acid and diluted to a concentration of 0.5 μg/μL. To generate the spectral library, a pooled sample (2 μg of each sample) was loaded onto an equilibrated, high-pH, reversed-phase fractionation spin column (84868, Thermo Fisher Scientific). A step gradient of 8 increasing acetonitrile concentrations (5, 7.5, 10, 12.5, 15, 17.5, 20 and 50% acetonitrile) in a volatile high-pH elution solution was then added to the columns to elute the peptides as eight different gradient fractions. The fractionated samples were then evaporated using vacuum evaporation and resuspended in 20 μL of 0.1% formic acid. Two microliters of each fraction was loaded for LC-DDA-MS/MS analysis.

### 6. LC-MS/MS analysis

The iRT reagent (Biognosys, Switzerland) was added at a ratio of 1:10 v/v to all peptide samples to calibrate the retention time of the extracted peptide peaks. For analysis, 1 μg of peptide from each sample was loaded into a trap column (75 μm * 2 cm, 3 μm, C18, 100 Å) at a flow rate of 0.4 μL/min and then separated with a reversed-phase analytical column (75 μm * 250 mm, 2 μm, C18, 100 Å). Peptides were eluted with a gradient of 4%-35% buffer B (0.1% formic acid in 80% acetonitrile) for 90 min and then analyzed with an Orbitrap Fusion Lumos Tribrid Mass Spectrometer (Thermo Fisher Scientific, Waltham, MA, USA). The LC settings were the same for both the DDA-MS and DIA-MS modes to maintain a stable retention time.

For the generation of the spectral library, the eight fractions obtained from the spin column separation was analyzed with mass spectrometry in DDA mode. The MS data were acquired in high-sensitivity mode. A full MS scan was acquired within a 350-1,500 m/z range with the resolution set to 120,000. The MS/MS scan was acquired in Orbitrap mode with a resolution of 30,000. The HCD collision energy was set to 30%. The AGC target was set to 5e4, and the maximum injection time was 45 ms.

The individual samples were analyzed in DIA-MS mode. The variable isolation window of the DIA method with 39 windows was used for DIA acquisition (Table S1). The full scan was obtained at a resolution of 60,000 with a m/z range from 350 to 1,400, and the DIA scan was obtained at a resolution of 30,000. The AGC target was 1e6, and the maximum injection time was 50 ms. The HCD collision energy was set to 32%. A single DIA analysis of pooled peptides was performed after every eight samples as the quality control.

### 7. Label-free DIA quantification

To generate a spectral library, ten DDA raw files were first searched by Proteome Discoverer (version 2.1; Thermo Scientific) with SEQUEST HT against the Swiss-Prot rat database (released in May 2019, containing 8,086 sequences). The iRT sequence was also added to the rat database. The search allowed two missed cleavage sites in trypsin digestion. Carbamidomethyl (C) was specified as the fixed modification. Oxidation (M) was specified as the variable modification. The parent ion mass tolerances were set to 10 ppm, and the fragment ion mass tolerance was set to 0.02 Da. The Q value (FDR) cutoff at the precursor and protein levels was 1%. Then, the search results were imported to Spectronaut Pulsar (Biognosys AG, Switzerland) software to generate the spectral library [25].

The QC and individual acquisition DIA files were imported into Spectronaut Pulsar with default settings. The peptide retention time was calibrated according to the iRT data. Cross-run normalization was performed to calibrate the systematic variance of the LC-MS performance, and local normalization based on local regression was used[26]. Protein inference was performed using the implemented IDPicker algorithm to generate the protein groups[27]. All results were then filtered according to a Q value less than 0.01 (corresponding to an FDR of 1%). The peptide intensity was calculated by summing the peak areas of the respective fragment ions for MS2. The protein intensity was calculated by summing the respective peptide intensity.

### 8. Statistical analysis

To analyze the proteomic data, we first used the sequential-KNN method to impute the missing value of QC samples[28]. The QC samples were used as technical replicates to evaluate the stability of mass spectrometry. Then, we required the proteins to be removed if the coefficient of variation (CV) value in the QC samples was larger than 0.3. The missing values in the remaining proteins of all the individual acquisition samples were then imputed using the sequential-KNN method.

Statistical analysis comparing three or four time points was performed by one-way ANOVA. The proteins identified on days 3, 6, 9 and 14 were compared with the control samples at the corresponding time points. The differential proteins were screened with the following criteria: proteins with at least two unique peptides were allowed; fold change ≥2 or ≤0.5; and P < 0.05 by independent sample t-test. Group differences resulting in P < 0.05 were identified as statistically significant. The P-values of group differences were also adjusted by the Benjamini and Hochberg method[29]. All results are expressed as the mean ± standard deviation.

### 9. Bioinformatics analysis

The differential proteins were analyzed by Gene Ontology (GO) based on biological processes, cellular components, and molecular functions using DAVID [30]. Protein interaction network analysis was performed using the STRING database (https://string-db.org/cgi/input.pl) and visualized by Cytoscape (V.3.7.1) [31] and OmicsBean workbench (http://www.omicsbean.cn). Biological pathway analysis and disease/biofunction analysis were performed by IPA software (Ingenuity Systems, Mountain View, CA, USA). The bubble figures of the three groups were visualized by the Wu Kong platform (https://www.omicsolution.org/wkomics/main/).

## Results

### 1. Characterization of simvastatin-induced rats

A total of 32 rats were subjected to simvastatin intragastric administration. The body weight of each rat was recorded for 14 days. Interestingly, we found that the simvastatin-treated rats exhibited different clinical manifestations. Some rats lost weight rapidly, while some of them seemed to have no change in body weight. Details of these 32 rats during the 14 days are presented in Table S2. Finally, nine rats survived for 14 days, five of whom lost weight significantly on day 4, while four of them did not show significant changes compared with the control group. In addition, another seven rats did not survive for 14 days due to excessive body weight loss (Figure 2A). Therefore, we divided these simvastatin-treated rats into three groups, namely, SAMS-non, SAMS and SAMS-severe.

**Figure 2.**
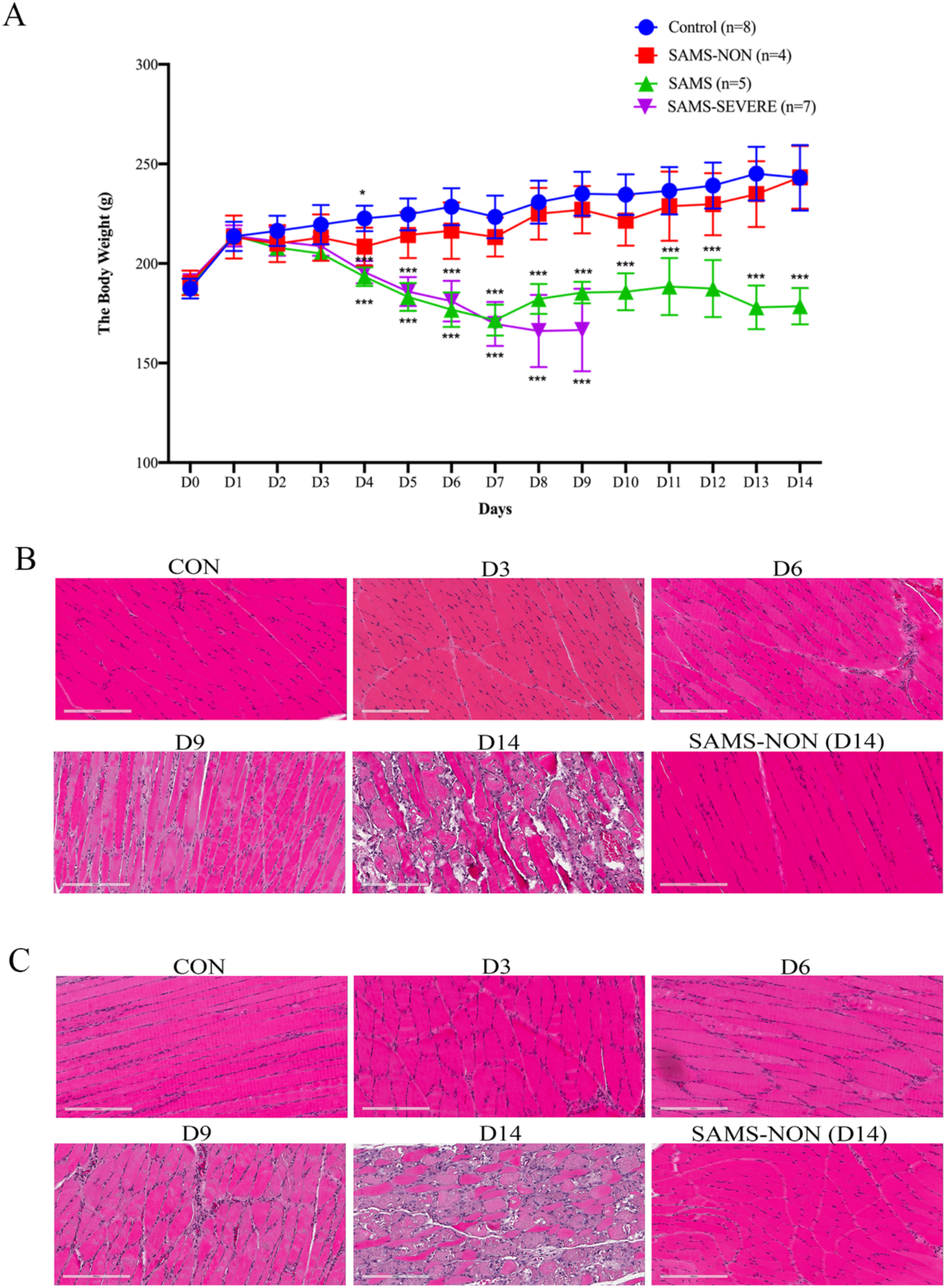
Body weight and pathological changes in the SAMS rat model. (A) Body weight changes of three groups in the SAMS rat model. The results are shown as the mean ± SD for the control group, the SAMS-non group, and the SAMS-severe group (\**P* < 0.05). (B). Periodic changes in gastrocnemius muscle on days 3, 6, 9 and 14 in the SAMS group and the SAMS-non group on day 14. (C). Periodic changes in the soleus muscle on days 3, 6, 9 and 14 in the SAMS group and the SAMS-non group on day 14.

**Figure 3.**
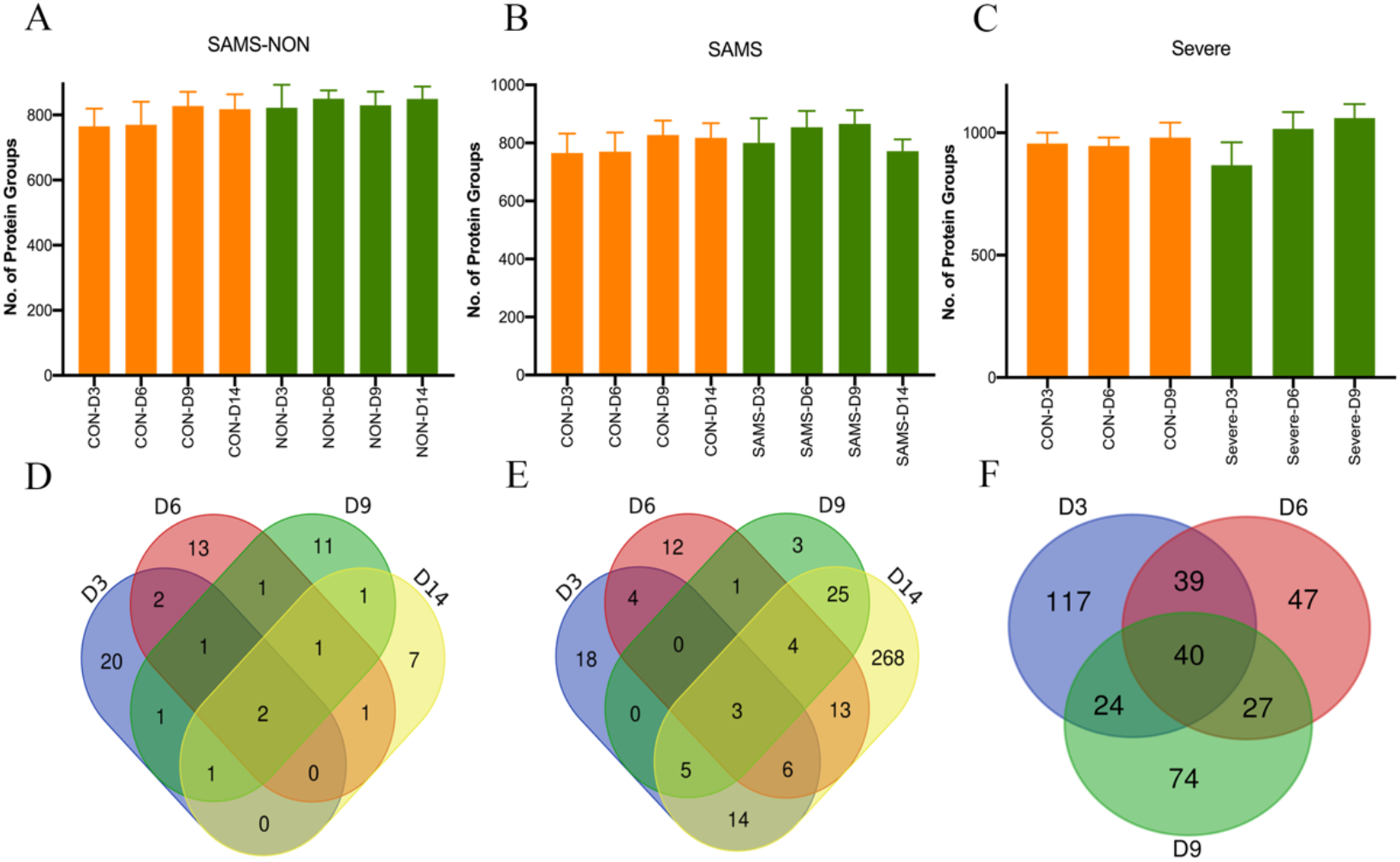
Proteomic analysis of urine samples of the three groups. (A-C) The distribution of quantified proteins in the SAMS-non group, SAMS group and SAMS-severe group. Error bars represent multiple independent samples. (D-E) Differential proteins at multiple time points in the SAMS-non group, SAMS group and SAMS-severe group.

The pathological changes in the simvastatin-treated rats at four time points are presented in Figure 2B, 2C. The gastrocnemius muscle and soleus muscle exhibited similar pathological progression during the 14 days. The interstitium of myocytes was enlarged, while the tissue morphology was normal on day 3. On day 6, a small number of inflammatory cells exhibited infiltration and hyperemia, and the tissue morphology remained normal. On day 9, the muscle fibers were clearly broken with obvious hyperemia and inflammatory cell infiltration, and some muscle fibers were lysed. On day 14, the muscle cells swelled with a large number of vacuoles, and the muscle fibers ruptured and dissolved severely. A large number of inflammatory cells exhibited infiltration and hyperemia with partial necrosis, which is very similar to rhabdomyolysis in humans. Overall, our research mimics the whole pathogenesis process of SAMS.

The biochemical parameter results are presented in Table 1. The concentrations of ALT, AST and CK were significantly elevated on day 3 but did not reach the 3× ULN. The concentration of BUN was significantly elevated on day 14, indicating possible kidney injury. The concentrations of CHO, TG, HDL, and LDL decreased on day 14.

**Table 1.** Biochemical index changes in the SAMS rat model.

|  | CON (n=4) | Day 3 (n=4) | Day 6 (n=4) | Day 9 (n=4) | Day 14 (n=4) | SAMS-non (n=4) |
| --- | --- | --- | --- | --- | --- | --- |
| ALT (U/L) | 25.33 ± 6.50 | 62.25 ± 20.18** | 84.50 ± 33.57** | 128.25 ± 86.70* | 123 ± 108.10 | 35.67 ± 14.52 |
| AST (U/L) | 58 ± 6.35 | 129 ± 38.54** | 244 ± 68.94** | 377.33 ± 372.50 | 415.55 ± 432.60 | 74.67 ± 10.34* |
| BUN (mmol/L) | 8.39 ± 1.43 | 6.8 ± 0.96 | 9.73 ± 2.78 | 9.36 ± 1.37 | 9.83 ± 3.01 | 8.60 ± 1.56 |
| CHO (mmol/L) | 1.49 ± 0.15 | 1.16 ± 0.20* | 1.56 ± 0.63 | 1.72 ± 0.04* | 1.55 ± 0.60 | 1.54 ± 0.22 |
| TG (mmol/L) | 0.40 ± 0.08 | 0.36 ± 0.06 | 0.55 ± 0.20 | 0.73 ± 0.33 | 0.43 ± 0.16 | 0.36 ± 0.07 |
| HDL (mmol/L) | 1.26 ± 0.12 | 0.72 ± 0.23** | 1.06 ± 0.50 | 1.30 ± 0.14 | 1.41 ± 0.50 | 1.34 ± 0.21 |
| LDL (mmol/L) | 0.17 ± 0.03 | 0.22 ± 0.02* | 0.20 ± 0.06 | 0.23 ± 0.03* | 0.20 ± 0.09 | 0.17 ± 0.03 |
| CK (U/L) | 184.33 ± 55.67 | 466.33 ± 135.95** | 968.25 ± 414.97** | 1926 ± 1783.33 | 1629.9 ± 2244.67 | 249 ± 35.69 |

Footnote: Values are given as the mean ± SD. Comparison between two groups were conducted using two-sided, unpaired t-test. (P\* < 0.05, P\*\* < 0.01).

The pathological changes and biochemical index parameter result of the remaining four SAMS-non and seven SAMS-severe rats are presented in Figure S1. For the SAMS-non group, the nuclei of the muscle cells remained normal, and there was no muscle fiber destruction and vacuole formation and no vascular congestion or inflammatory cell infiltration in the gastrocnemius and soleus muscles, which was very similar to the control group. For the SAMS-severe group, the pathological changes were very similar to those of the SAMS group on day 14. According to these above results, we believed that day 3 was the early time point of SAMS.

### 2. Urine proteome changes in simvastatin-induced rats

We first analyzed the urine proteome of the SAMS (n=44) and SAMS-non groups (n=40) using label-free DIA-LC-MS/MS quantitation. These two groups had the same control group. Then, we also analyzed the urine proteome of the SAMS-severe group (n=37). A total of 810 urinary proteins with at least 2 unique peptides were identified with <1% FDR at the protein level in the SAMS and SAMS-non groups, while a total of 972 urinary proteins with at least 2 unique peptides were identified in the SAMS-severe group.

We first evaluated the quality control of the proteomic data. The median technical CV in the QC samples of the SAMS and SAMS-non groups was 0.17, while in the SAMS-severe group, the median technical CV was 0.16, indicating the great stability of mass spectrometry in our research (Figure S2A, S2B). In addition, the line correlation of the QC samples also indicates the great stability of mass spectrometry in this research (Figure S2C, S2D).

After screening the proteins in the QC samples with a CV less than 0.3, a total of 680 proteins were considered to be highly specific to the SAMS and SAMS-non groups, while a total of 825 proteins were considered to be highly specific to the SAMS-severe group. Then, the missing values of proteomic data biological samples in the SAMS, SAMS-non and SAMS-severe groups were imputed with the sequential-KNN method. Finally, 628 proteins were retained in the SAMS and SAMS-non groups, while 758 proteins were retained in the SAMS-severe group for further differential urine protein selection. The distribution of quantified proteins in the SAMS-non group, SAMS group and SAMS-severe group at different time points is presented in Figure 2A-C.

After using the screening criteria, 49, 42, 40 and 337 differential proteins were significantly changed in the SAMS group, while 27, 21, 19 and 13 differential proteins were significantly changed in the SAMS-non group on days 3, 6, 9, and 14, respectively. For the SAMS-severe group, 220, 153 and 165 differentially expressed proteins were significantly changed on days 3, 6, and 9 (fold change ≥2 or ≤ 0.5, P < 0.05, Table S3). The Venn diagram of these three groups is presented in Figure 2D-2E. Most of the differential proteins changed uniquely at these three or four time points, indicating that different biological changes may occur at different time points in SAMS pathogenesis.

### 3. Functional analysis of the differential proteins in simvastatin-induced rats

As the differential proteins in these three groups at different time points varied greatly, we supposed that the biological functions of these three groups at different time points were different. Therefore, we mainly focused on the biological processes, pathways and disease/biofunction changes on day 3 in these three groups. The functional annotation of the SAMS group on day 14 is also presented in this section.

We noticed that the majority of the biological processes in the three groups were different starting on day 3 (Figure 4A). The SAMS-non group was more associated with the response to nutrient levels, phospholipid efflux, innate immune response in mucosa, leukocyte cell-cell adhesion, lipoprotein metabolic process, cholesterol efflux and gluconeogenesis. The SAMS group exhibited more association with the negative regulation of endopeptidase activity, acute-phase response, receptor-mediated endocytosis, cellular response to interleukin-6, complement activation (classical pathway), vitamin metabolic process, hemoglobin import and carbohydrate catabolic process. Interestingly, in the SAMS-severe group, differential proteins were more associated with cell adhesion, aging, glutathione metabolic process, complement activation (classical pathway), acute-phase response, blood coagulation, response to cytokines and oxidation-reduction process. We also found that for the pathways (Figure 4B, 4C) and the disease/biofunctions (Figure 4D, 4E) on day 3, the SAMS-non group and the SAMS group exhibited associations with acute phase response signaling, the coagulation system, clathrin-mediated endocytosis signaling, glucocorticoid receptor signaling, degranulation of blood platelets, aggregation of cells, inflammation of organs, adhesion of immune cells, metabolism of cellular proteins, chronic inflammatory disorders, coagulation of blood, glucose metabolism disorders and aggregation of blood platelets. Interestingly, we noticed that the production of nitric oxide and reactive oxygen species in macrophages, necrosis and bleeding were only present in the SAMS-non group compared with the SAMS group. Some biological functions showed positive or negative regulation in the SAMS-severe group. For example, IL-15 production, NF-κB signaling, IL-6 signaling, sirtuin signaling pathway, cell survival, adhesion of blood cells, cell movement of smooth muscle cells, adhesion of immune cells, and cell movement of muscle cells were upregulated, while the PPAR signaling, xenobiotic metabolism general signaling pathway, glutathione redox reactions I, bleeding, apoptosis, and necrosis were downregulated.

**Figure 4.**
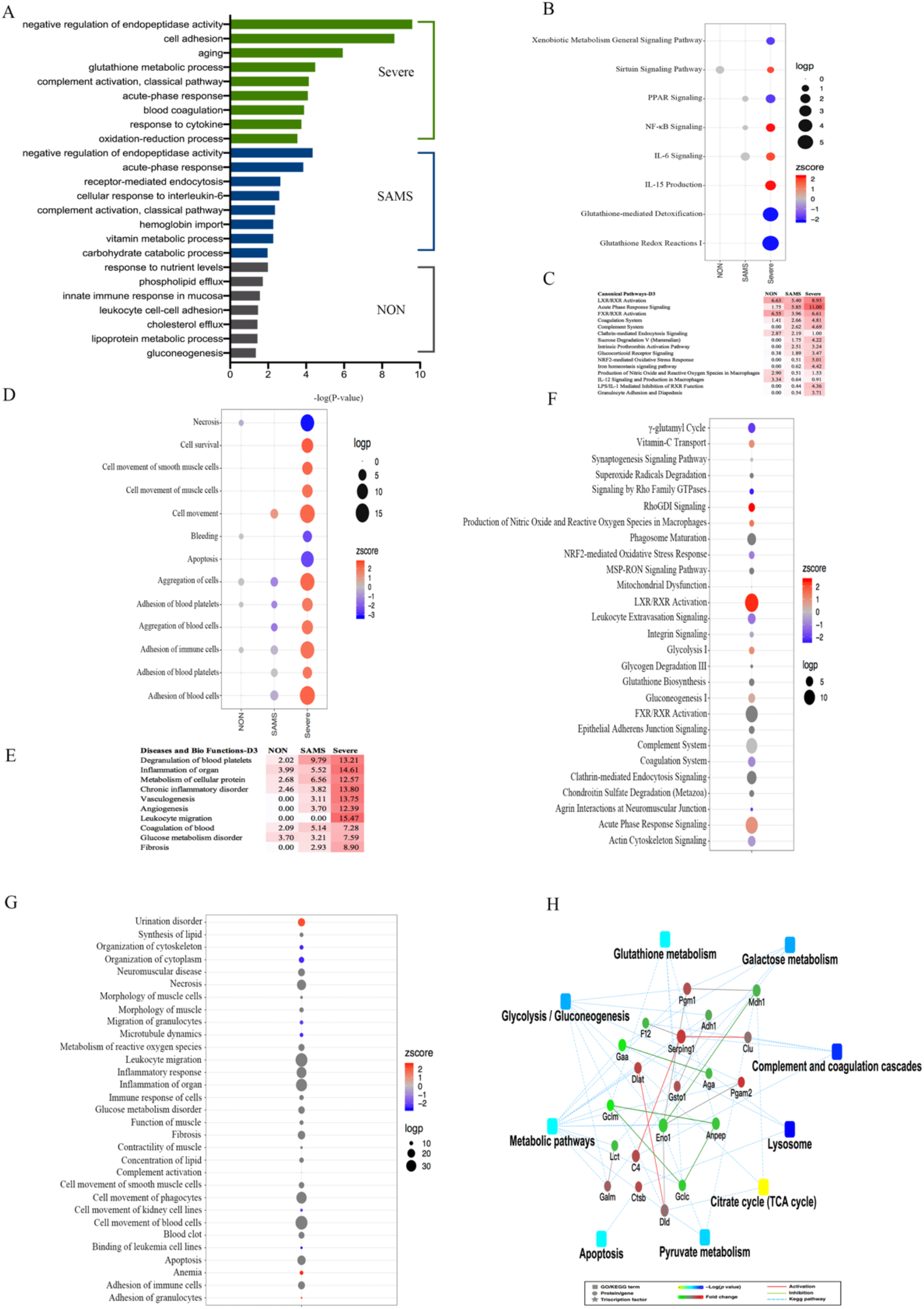
Functional analysis of differential proteins in the SAMS-non, SAMS and SAMS-severe groups. (A) The top 10 biological processes of differentially expressed proteins in the SAMS-non, SAMS and SAMS-severe groups on day 3. (B and D) Representative pathways of differential proteins in the SAMS-non, SAMS and SAMS-severe groups on day 3. The z-score algorithm was used to predict the activation state (either activated or inhibited) of the pathways. If the z-score ≤ −2, the pathway is predicted to be statistically significantly inhibited. (C and E) Representative disease and biofunction of differential proteins in the SAMS-non, SAMS and SAMS-severe groups on day 3. The z-score algorithm was used to predict the activation state (either activated or inhibited) of the disease and biofunction. If the z-score ≤ −2, the disease and biofunction list is predicted to be statistically significantly inhibited. (F) Representative pathways of differential proteins in the SAMS group on day 14. (G) Representative disease and biofunction of differential proteins in the SAMS group on day 14. (H) The interaction diagram of proteins of representative KEGG pathways. Green solid lines represent inhibition; red solid lines represent activation. The color bar from red to green represents the fold change of the protein level from increasing to decreasing. The significance of the pathways represented by −log(p-value) is shown by color scales with dark blue as the most significant.

On day 14 in the SAMS group, we found that the LXR/RXR activation, RhoGDI signaling, urination disorder, anemia, adhesion of granulocytes and complement activation were upregulated, while the γ-glutamyl cycle, signaling by Rho family GTPases, agrin interactions at neuromuscular junctions, organization of cytoplasm, organization of cytoskeleton, migration of granulocytes, microtubule dynamics, cell movement of kidney cell lines and binding of leukemia cell lines were downregulated. In addition, we also noticed that pathways and disease/biofunctions such as complement system, clathrin-mediated endocytosis signaling, leukocyte extravasation signaling, coagulation system, gluconeogenesis I, vitamin-C transport, NRF2-mediated oxidative stress response, glycolysis I, cell movement of blood cells, inflammation of organ, inflammatory response, apoptosis, neuromuscular disease, glucose metabolism disorder, cell movement of smooth muscle cells, concentration of lipid, morphology of muscle and synthesis of lipid were also enriched on day 14 (Figure 4F, 4G).

Differential proteins such as AGA, CTSB, F12, CLU, C4, SERPING1, PGM1, GAA, ADH1, GALM, PGAM2, DID, PGM1, ENO1, DIAT, MDH1, GCLM, GCLC and ANPEP were involved in KEGG pathways such as lysosome, complement and coagulation cascade, galactose metabolism, glycolysis/gluconeogenesis, pyruvate metabolism, citrate cycle (TCA cycle) and glutathione metabolism (Figure 4H).

### 4. Potential biomarkers to predict muscle damage in different phases

We compared the differential proteins on day 3 in these three groups and found that more than half of the DEPs were unique in these three groups, indicating different disease progression beginning on day 3 (Figure 5A). We then focused on the human homologous proteins on day 3 in the SAMS-non and SAMS groups. For the SAMS-non group, the protein panel including AT1A1, S12A3, UFD1, NGAL, FETUB, UROM, GPDA, PGAM1, DEFB1, VDAC1, HMGB2, LYSC1, ITPA, VTDB, APOA4 and HEP2 indicated the possibility that SAMS may not develop. For the SAMS group, the protein panel including PLBL1, S10AB, FIBG, CFAI, EHD1, OSTP, UBB, AMPE, HEXA, CUBN, FINC, CRP, ANTR1 and DHSO was able to indicate the early phase of SAMS before there was an increase in CK to three times the upper limit of normal. Among these 14 differential proteins, FIBG, OSTP and CRP were reported to be associated with skeletal muscle damage, while EHD1, CUBN and FINC were reported to be associated with the mechanism of SAMS (Table 2).

**Figure 5.**
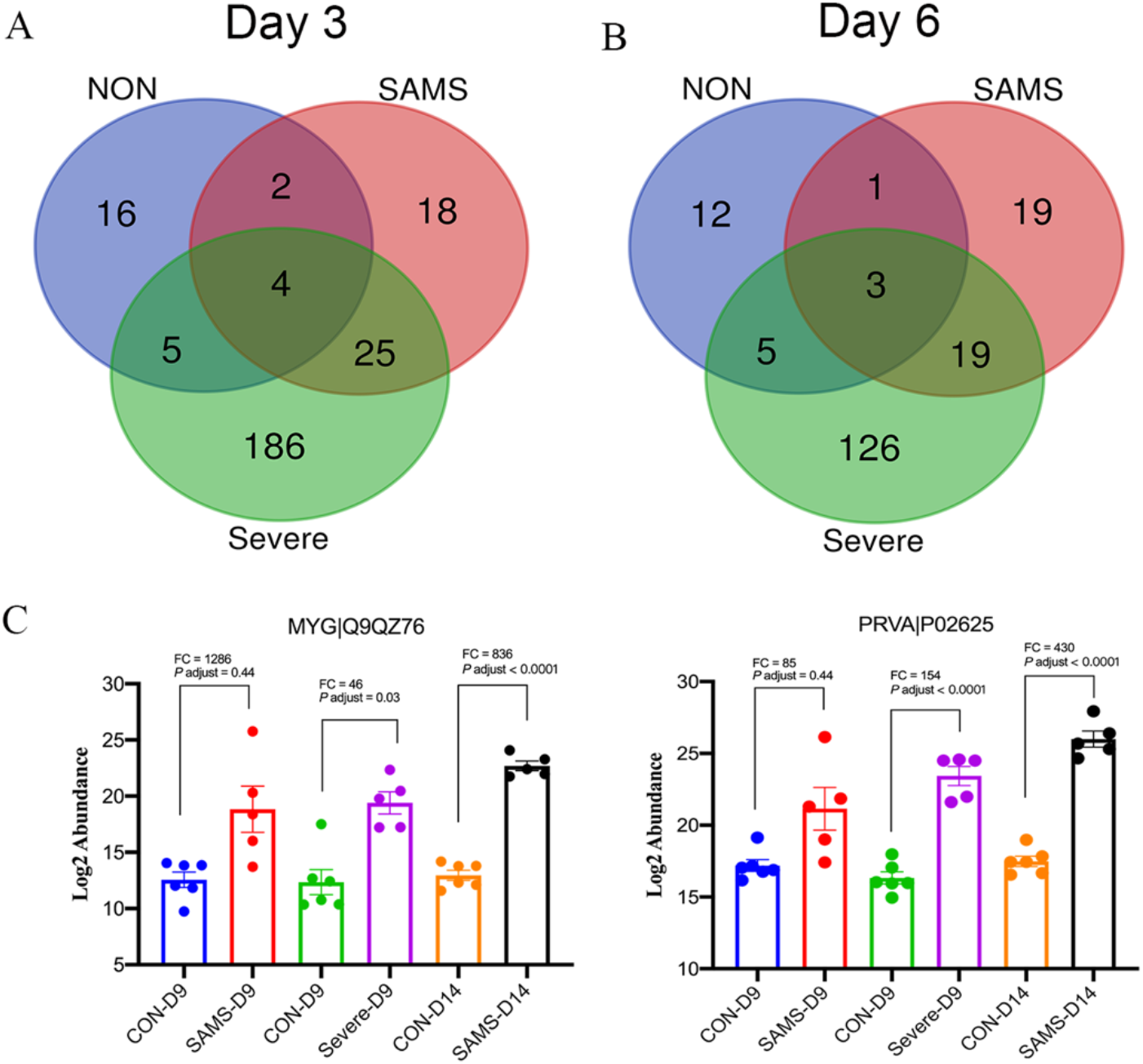
Potential biomarkers to predict the extent of muscle damage. (A) Venn diagram of differential proteins in the SAMS-non, SAMS and SAMS-severe groups on day 3. (B) Venn diagram of differential proteins in the SAMS-non, SAMS and SAMS-severe groups on day 6. (C) Scatter plot graphs showing two proteins that are potential biomarkers to predict severe muscle damage. The x-axis shows the different stages of muscle damage. The y-axis shows the log2 of the normalized abundance based on DIA quantification. The fold change was calculated based on the average group of normalized abundances. Data are presented as the mean ± SEM.

**Table 2.** Potential biomarkers to predict early muscle damage.

| UniProt ID | UniProt accession | Human ortholog | Protein name | Trend | P-value | Fold change | Related to muscle damage | Related to SAMS mechanism |
| --- | --- | --- | --- | --- | --- | --- | --- | --- |
| PLBL1_RAT | Q5U2V4 | Q6P4A8 | Phospholipase B-like 1 | ↑ | 3.81E-02 | 8.58 | - | - |
| S10AB_RAT | Q6B345 | P31949 | Protein S100-A11 | ↑ | 1.36E-02 | 7.64 | - | - |
| FIBG_RAT | P02680 | P02679 | Fibrinogen gamma chain | ↑ | 2.57E-02 | 6.36 | [32] | - |
| CFAI_RAT | Q9WUW3 | P05156 | Complement factor I | ↑ | 2.89E-02 | 3.87 | - | - |
| EHD1_RAT | Q641Z6 | Q9H4M9 | EH domain-containing protein 1 | ↑ | 4.83E-03 | 2.48 | - | [33] |
| OSTP_RAT | P08721 | P10451 | Osteopontin | ↑ | 1.33E-02 | 2.35 | [34] | - |
| UBB_RAT | P0CG51 | P0CG47 | Polyubiquitin-B | ↑ | 4.75E-02 | 2.10 | - | - |
| AMPE_RAT | P50123 | Q07075 | Glutamyl aminopeptidase | ↓ | 2.22E-02 | 0.50 | - | - |
| HEXA_RAT | Q641X3 | P06865 | Beta-hexosaminidase subunit alpha | ↓ | 4.49E-02 | 0.45 | - | - |
| CUBN_RAT | O70244 | O60494 | Cubilin | ↓ | 6.34E-03 | 0.43 | - | [35] |
| FINC_RAT | P04937 | P02751 | Fibronectin | ↓ | 1.77E-04 | 0.42 | - | [36] |
| CRP_RAT | P48199 | P02741 | C-reactive protein | ↓ | 3.35E-02 | 0.40 | [37] | - |
| ANTR1_RAT | Q0PMD2 | Q9H6X2 | Anthrax toxin receptor 1 | ↓ | 2.83E-03 | 0.39 | - | - |
| DHSO_RAT | P27867 | Q00796 | Sorbitol dehydrogenase | ↓ | 4.66E-02 | 0.38 | - | - |

We also compared the differential proteins in the monitoring phase (day 6) of these three groups. For the SAMS group, 19 unique urinary proteins changed significantly on day 6 (Figure 5B). Details of 17 human homologous proteins are presented in Table 3. Among these 17 differential proteins, APOC1 and LDLR were reported to be associated with skeletal muscle damage, while FINC and LDLR were reported to be associated with the mechanism of SAMS (Table 3).

**Table 3.** Potential biomarkers to monitor muscle damage.

| UniProt ID | UniProt accession | Human ortholog | Protein name | Trend | P-value | Fold change | Related to muscle damage | Related to SAMS mechanism |
| --- | --- | --- | --- | --- | --- | --- | --- | --- |
| ACPH_RAT | P13676 | P13798 | Acylamino-acid-releasing enzyme | ↑ | 4.39E-03 | 5.35 | - | - |
| MUCEN_RAT | Q6AY82 | Q9ULC0 | Endomucin | ↑ | 4.65E-02 | 4.59 | - | - |
| DPEP1_RAT | P31430 | P16444 | Dipeptidase 1 | ↑ | 3.22E-02 | 4.22 | - | - |
| ACY2_RAT | Q9R1T5 | P45381 | Aspartoacylase | ↑ | 3.68E-02 | 3.13 | - | - |
| ABHEB_RAT | Q6DGG1 | Q96IU4 | Protein ABHD14B | ↑ | 1.32E-02 | 2.52 | - | - |
| S10A6_RAT | P05964 | P06703 | Protein S100-A6 | ↑ | 3.17E-02 | 2.42 | - | - |
| GALM_RAT | Q66HG4 | Q96C23 | Aldose 1-epimerase | ↑ | 1.67E-02 | 2.01 | - | - |
| FINC_RAT | P04937 | P02751 | Fibronectin | ↓ | 2.15E-02 | 0.50 | - | [36] |
| TKFC_RAT | Q4KLZ6 | Q3LXA3 | Triokinase/FMN cyclase | ↓ | 4.61E-02 | 0.49 | - | - |
| ATL4_RAT | Q4FZU4 | Q6UY14 | ADAMTS-like protein 4 | ↓ | 4.03E-02 | 0.37 | - | - |
| AMBP_RAT | Q64240 | P02760 | Protein AMBP | ↓ | 3.25E-02 | 0.32 | - | - |
| APOC1_RAT | P19939 | P02654 | Apolipoprotein C-I | ↓ | 1.16E-02 | 0.32 | [38] | - |
| PPT1_RAT | P45479 | P50897 | Palmitoyl-protein thioesterase 1 | ↓ | 6.02E-06 | 0.30 | - | - |
| CCN3_RAT | Q9QZQ5 | P48745 | CCN family member 3 | ↓ | 1.43E-02 | 0.27 | - | - |
| NID2_RAT | B5DFC9 | Q14112 | Nidogen-2 | ↓ | 4.22E-02 | 0.26 | - | - |
| LDLR_RAT | P35952 | P01130 | Low-density lipoprotein receptor | ↓ | 2.48E-02 | 0.25 | [39] | [40] |
| VAS1_RAT | O54715 | Q15904 | V-type proton ATPase subunit S1 | ↓ | 6.30E-04 | 0.17 | - | - |

Notably, we also found that myoglobin (MYG) and parvalbumin alpha (PRVA) increased dramatically in the late phase on days 9 and 14 of the SAMS group and on day 9 of the SAMS-severe group (Figure 5B). As rhabdomyolysis is characterized by muscle necrosis, which causes the release of myoglobin into the bloodstream, we therefore suppose that these two proteins may combine with CK to predict the extent of muscle injury, as they are more sensitive than CK elevation. The downregulation of these two proteins may predict the recovery of muscle damage in the future.

### 5. Protein-protein interaction network analysis

To better understand the pathogenic mechanism of SAMS, the PPI network was generated by the STRING database based on differential proteins of the SAMS group on day 14. The PPI enrichment p-value was less than 1.0e-16, indicating that these DEPs have more interactions among themselves (Figure S3). Then, we used cytoHubba to predict the top 15 key hub proteins and generated the subnetwork based on the node degree. C3, APP, KNG1, APOE, FGG, P4HB, EGF, CST3, SPP1, AGT, GC, SPERIND1 and CDH2 exhibited strong associations with each other, indicating that these proteins may play an important role in the development of SAMS (Figure 6A). In addition, we also generated the protein-protein interactions involved in the important GO terms (Figure 6B). Differential proteins such as ACE, LGMN, CRP, GCLC, CRK, B2M, PRDX3, C4BPA, GSN, C4BPB, CLU, ENO1, KNG1, P4HB, C4, CFD, C5, SOD1, MAP1, SERPING1, GGT1 and AGT were involved in biological processes such as proteolysis, metabolic processes, response to stress, response to reactive oxygen species, regulation of immune system processes, inflammatory response, catabolic processes and response to oxidative stress.

**Figure 6.**
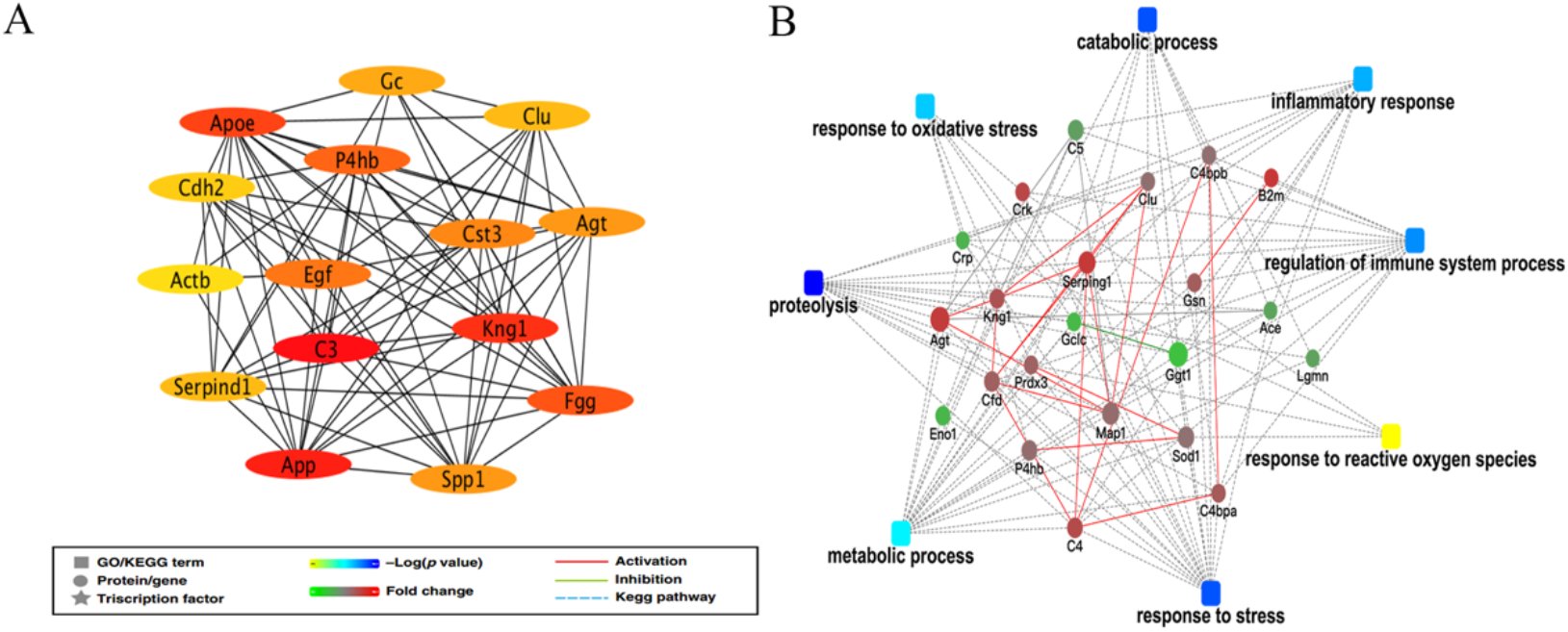
Interaction network analysis of differential proteins in the SAMS group on day 14. (A) The top 15 key hub protein interaction networks of differential proteins in the SAMS group on day 14. (B) The interaction diagram of proteins of representative GO terms. Green solid lines represent inhibition; red solid lines represent activation. The color bar from red to green represents the fold change of the protein level from increasing to decreasing. The significance of the GO terms represented by −log(p-value) is shown by color scales with dark blue as the most significant.

## Discussion

The urine proteome has been applied in various aspects, such as early disease detection[41], phenotyping[42] and diagnosis[43,44]. However, the urine proteome has not been applied to investigate the possible side effects of drugs. In this study, we established a SAMS rat model to identify urine proteome changes associated with simvastatin-induced muscle damage. Three different clinical manifestations (SAMS-non, SAMS and SAMS-severe) appeared throughout the entire course of this research. We therefore sought to detect different urine proteome changes in these three different clinical manifestations. Specifically, we focused on early urine proteome changes on day 3 in these three groups and urine proteome changes on day 14 in the SAMS group.

We found that some biological functions on day 14 in the SAMS group were associated with the pathological mechanism of SAMS. For example, i) Rho family GTPase signaling was downregulated in the SAMS group on day 14. As Rho proteins are regarded as substrates for protein isoprenylation[45] and isoprenylation is a posttranslational modification requiring intermediates of the cholesterol biosynthesis pathway, we therefore suppose that downregulated Rho family GTPase signaling may inhibit the synthesis of cholesterol[46]. ii) Glycolysis/gluconeogenesis, citrate cycle (TCA cycle), pyruvate metabolism, glucose metabolism disorder and glycogen degradation III were all associated with mitochondrial dysfunction, while mitochondrial dysfunction has been reported to play an important role in the myotoxicity of statins in many studies[47–50]. iii) The NRF2-mediated oxidative stress response, production of nitric oxide and reactive oxygen species in macrophages, superoxide radical degradation and metabolism of reactive oxygen species were associated with oxidative stress. Mitochondria are the most important producers of reactive oxygen species (ROS) in cells[51], and simvastatin increases mitochondrial superoxide production in primary human skeletal muscle cells[49]. As hydrogen peroxide (H2O2) can be reduced to water by glutathione peroxidases, impairment in the antioxidative defense system could therefore be a susceptibility factor in patients developing statin-induced myotoxicity[52]. Interestingly, in our research, we also found that the γ-glutamyl cycle was downregulated in the SAMS group on day 14, while glutathione biosynthesis and glutathione metabolism were also enriched. iv) Finally, apoptosis was also enriched in the SAMS group. It has been reported that mitochondria play an important role in initiating the intrinsic pathway of apoptosis[53], and induction of apoptosis by statins has been demonstrated in human skeletal muscle cell lines[54] and in primary human myotubes[49]. These findings suggest that mitochondrial ROS production and apoptosis are closely linked. In addition to the above findings, we also found that the concentration of lipids and the synthesis of lipids were enriched, which is consistent with statins being used as lipid-lowering agents. The coagulation system was downregulated slightly (z-score: −0.816), which is consistent with the pharmacological effect of statins[55]. Complement activation was upregulated, and the inflammatory response was also enriched, indicating that this may be associated with immune-mediated necrotizing myopathy (IMNM)[56].

Interestingly, we noticed that various biological functions on day 3 in the SAMS-severe group were similar to those in the SAMS group on day 14. For example, glutathione redox reaction I and glutathione-mediated detoxification were also downregulated, and the oxidation-reduction process and glutathione metabolic process were also enriched. To our surprise, we found that the adhesion of blood platelets and the aggregation of blood platelets were upregulated in the SAMS-severe group, while the aggregation of blood platelets was downregulated in the SAMS group on day 3. As bleeding was also downregulated in the SAMS-severe group starting on day 3, we therefore suppose that this may be due to liver damage in the SAMS-severe group, but this damage did not appear in the SAMS group on day 3. When comparing biological functions between the SAMS and SAMS-non groups on day 3, we noticed that the complement system and the intrinsic prothrombin activation pathway were specifically enriched. In the SAMS-non group, the response to nutrient levels, phospholipid efflux, cholesterol efflux and lipoprotein metabolic processes were specifically enriched, which was associated with the pharmacological effect of statins as lipid-lowering agents. Taken together, we suppose that the SAMS-non group may be associated with the pharmacological effect of statins, that the SAMS group reflects the whole pathological process of SAMS, while the SAMS-severe group reflects muscle toxicity beginning on day 3 in this research.

Three differential proteins in the SAMS group had been reported to be associated with muscle damage since day 3. For example, i) the fibrinogen gamma chain (FIBG) increased ≥10-fold after impact trauma in the muscle tissue of male Wistar rats[32]; ii) osteopontin (OSTP) was reported to be increased in patients with idiopathic inflammatory myopathies, both in muscle and in serum[34]; and iii) C-reactive protein (CRP) was reported to be elevated only in humans with high CK levels when undergoing muscle-damaging exercise [37]. In addition, five differential proteins were associated with the mechanism of SAMS. For example, i) EH domain-containing protein 1 (EHD1) was reported to be required for perinuclear localization of GLUT4, which is regarded as the glucose transporter in muscle tissues[33]. ii) It has been reported that the main circulating metabolite of vitamin D accumulates in skeletal muscle cells, which is mediated by muscle cell uptake of circulating vitamin D-binding protein (DBP) through a megalin-cubilin membrane transport process[35]. Therefore, downregulated cubilin (CUBN) may affect vitamin D synthesis in muscle cells. iii) Last, fibronectin (FINC) levels were substantially reduced in the aged stem cell niche in skeletal muscle, leading to detrimental consequences for the function and maintenance of muscle stem cells [36].

Nonetheless, several limitations of the study should be considered. First, our research ended on day 14, and it is unclear whether the SAMS-non group will develop SAMS at later stages. Second, given the relatively small number of investigated rats in this preliminary study, we are well aware that our results still need clinical validation in future studies. A larger number of clinical urine samples is needed to validate the differential proteins in our research to validate their sensitivity and specificity.

## Conclusions

In summary, we aimed to observe urine proteome changes associated with SAMS. Our results indicated that the urine proteome can reflect early changes in the SAMS rat model, providing the potential for monitoring drug side effects in clinical research.

## Conflict of interest/disclosure statement

The authors declare no competing financial interests.

## Acknowledgments

This work was supported by the National Key Research and Development Program of China (2018YFC0910202 and 2016YFC1306300); the Fundamental Research Funds for the Central Universities (2020KJZX002); the Beijing Natural Science Foundation (7172076); the Beijing Cooperative Construction Project (110651103); the Beijing Normal University (11100704); and the Peking Union Medical College Hospital (2016-2.27).

## Supplemental Materials

**Table S1.** The variable isolation window information of the DIA method with 36 windows. (A) (B)

**Table S2.** Details of the 32 rats during the 14-day treatment with simvastatin.

**Table S3.** Differential proteins identified in the three groups. (A) Differential proteins identified on days 3, 6, 9, and 14 in the SAMS-non group. (B) Differential proteins identified on days 3, 6, 9, and 14 in the SAMS group. (C) Differential proteins identified on days 3, 6 and 9 in the SAMS-severe group.

**Figure S1.** The pathological changes and biochemical index parameters of the remaining four SAMS-non and seven SAMS-severe rats.

**Figure S2.** Quality control of the proteomic data. (A) CV distribution of the QC samples in the SAMS-non and SAMS groups. (B) CV distribution of the QC samples in the SAMS-severe group. (C) Linear correlation of the QC samples in the SAMS-non and SAMS groups. (D) Linear correlation of the QC samples in the SAMS-severe group.

**Figure S3.** The PPI network of differential proteins identified on day 14 in the SAMS group.

## Notes

### Competing Interest Statement

The authors have declared no competing interest.

